# Canagliflozin Extends Lifespan in Genetically Heterogeneous Male But Not Female Mice

**DOI:** 10.1101/2020.05.23.112763

**Authors:** Richard A. Miller, David E. Harrison, David B. Allison, Molly Bogue, Vivian Diaz, Elizabeth Fernandez, Andrzej Galecki, W. Timothy Garvey, Navasuja Kumar, Martin A. Javors, Warren C. Ladiges, Francesca Macchiarini, James Nelson, Peter Reifsnyder, Nadia A. Rosenthal, Adam B. Salmon, Daniel L. Smith, Jessica M. Snyder, David B. Lombard, Randy Strong

## Abstract

Canagliflozin (Cana) is an inhibitor of the sodium glucose transporter 2 (SGLT2), and is thought to act by blocking renal reuptake and intestinal absorption of glucose. Cana is FDA-approved for treatment of diabetes, and affords protection from cardiovascular and kidney diseases. In the context of the mouse Interventions Testing Program, genetically heterogeneous mice were given chow containing 180 ppm Cana at 7 months of age until their death. Cana extended median survival of male mice by 14%, with p < 0.001 by log-rank test. Cana also increased by 9% the age for 90^th^ percentile survival (p < 0.001 by Wang/Allison test), with parallel effects seen at each of three test sites. Cana did not alter the distribution of inferred cause of death, nor of incidental pathology findings at end-of-life necropsies. No benefits were seen in female mice. The lifespan benefit of Cana is likely to reflect blunting of peak glucose levels, because similar longevity effects are seen in mice given acarbose, a diabetes drug that blocks glucose surges through a distinct mechanism, i.e. slowing breakdown of carbohydrate in the intestine. Interventions that control daily peak glucose levels deserve attention as possible preventive medicines to protect from a wide range of late-life neoplastic and degenerative diseases.

## Introduction

Aging is the dominant risk factor for most chronic diseases that afflict people in industrialized societies. There is now ample evidence, in mice, that the process of aging can be delayed or retarded by low-calorie diets (1, 2), by natural or engineered mutations in any of several genes that modify growth hormone and IGF-1 signals (3, 4), and, more recently, by drugs added to food (5). Each of these approaches has been shown to extend lifespan and also to delay multiple forms of late-life illness, including both neoplastic and degenerative diseases. Identification of new drugs that extend mammalian lifespan is of great interest from at least two perspectives. First, elucidating the mechanism of action of such agents will provide new insights into the biology of the aging process and into the links between aging and multiple forms of late-life diseases. Second, these drugs may provide a starting point for development of anti-aging interventions in humans.

The Interventions Testing Program (ITP), supported by the National Institute on Aging, tests the effects of drugs on mouse lifespan (http://www.nia.nih.gov/research/dab/interventions-testing-program-itp)(6, 7). ITP studies use annual cohorts of genetically heterogeneous mice (UM-HET3) equally distributed among three testing sites. Six compounds have thus far produced significant extension of lifespan in ITP studies in one or both sexes: rapamycin (8-10), 17α-estradiol (17aE2)(11, 12), nordihydroguaiaretic acid (NDGA) (12, 13), Protandim (11), glycine (14) and acarbose (12, 15). A seventh agent, aspirin, produced significant lifespan extension, in males, in an early study (13), but failed to show a benefit at two higher doses in a follow-up study (unpublished data).

The ITP experiment with Cana was motivated in part by our results using acarbose, a drug used in the treatment of type 2 diabetes (T2DM). Acarbose functions by inhibiting the breakdown of complex carbohydrates in the small intestine, thereby slowing the pace of glucose uptake and blunting the normal post-prandial spike in blood glucose levels. In humans, acarbose treatment reduces glycated hemoglobin (HbA1c) levels (16), although this effect was not seen in the ITP acarbose-treated mice (12). Acarbose, started at 4 months of age, led to a 22% increase in median lifespan in UM-HET3 males, and a significant but much smaller, 5%, increase in females (12). A test for effects on maximum lifespan (proportion alive at the 90^th^ percentile) was significant in both sexes. When initiated at 16 months, acarbose leads to significant lifespan extension in males but a smaller effect in females (11). Acarbose-treated male mice also show a delay in age-dependent pathology in multiple systems, including inflammatory changes in the hypothalamus (17). Taken together, these results suggest that blunting of post-prandial glucose peaks could delay aging and its consequences, including the illnesses that lead to death in mice, but they do not exclude other possible mechanisms for acarbose action.

These acarbose results prompted us to consider whether other drugs that inhibit surges in post-prandial blood glucose levels might also exert beneficial effects on longevity. Canagliflozin (Cana) is an FDA-approved diabetes drug that acts primarily by inhibition of the sodium glucose transporter 2 (SGLT2) in the proximal tubule of the kidney (18). In non-diabetic individuals, SGLT2 is responsible for up to 97% of the reabsorption of the 160-180 g of glucose filtered by the kidney per day. To a lesser extent, Cana also inhibits SGLT1, a related glucose transporter with broader distribution, expressed in the small intestine, kidney, and heart, among other tissues. In a large group of patients with T2DM and elevated cardiovascular risk, Cana reduced death from cardiovascular causes, as well as non-fatal strokes, myocardial infarction, and heart failure (19). In T2DM patients with kidney disease, Cana greatly reduced the risk of progression to kidney failure, as well as the risk of cardiovascular events. Cana, like other SGLT2 inhibitors, reduces fasting blood glucose, HbA1c levels, and body weight, while modestly increasing blood ketone levels (20). Like acarbose, Cana suppresses the post-prandial glucose surge (21, 22). The cardioprotective effects of SGLT2 inhibitors may be independent, at least partly, of their glycemic effects (23). In view of our previous results using acarbose, and the evidence for beneficial effects of SGLT2 inhibitors in humans, we conducted a test of the hypothesis that Cana would extend mouse lifespan.

## Methods

### Mice

These experiments used mice of the UM-HET3 genetically heterogeneous stock, produced by mating CByB6F1 mothers to C3D2F1 fathers. Each HET3 mouse is genetically unique, but each shares 50% of its genetic endowment with every other HET3 mouse, and the population thus consists of full sibs with respect to nuclear genes. Mice from second and subsequent litters were weaned into cages containing 3 male or 4 female mice, and assigned to Control or treatment (Cana) group using a random number table. Mice at each test site were produced over a period of six months in 2016 in roughly equal monthly cohorts. The C2016 cohort also included mice treated with other agents, and the results of those studies will be reported separately. Mice in the Cana group received this agent at 180 mg per kg of chow, from 7 months of age; this dose is equivalent to 30 milligrams per kg mouse body weight for a 30-gram mouse eating 5 grams of food each day. Mice were weighed at 6, 12, 18, and 24 months, but not subjected to any other manipulation. Cages were inspected daily, and deaths noted. Mice which were deemed unlikely to live for more than another 24 hr, based on a symptom checklist, were euthanized for humane reasons, with the day of euthanasia taken as the best estimate of the date of natural death for statistical purposes. Mice removed from the study, either for fighting or for other technical reasons (e.g., chip ID dysfunction, escape, accidental injury), are treated as known to be alive on the day of removal but lost to follow□up at that point. These “removed” mice are included in the Kaplan-Meier survival calculations, but not in calculations of median or 90^th^ percentile survival age. Removed mice made up 4.8% of the original Control group, and 3.8% of the Cana group. Details of husbandry conditions, such as cage changes, base diet, temperature, and light/dark cycle have been described extensively in other ITP reports (8, 11). Specific-pathogen free status is assessed quarterly at each site using a combination of serological and molecular methods, and all such tests have been negative for each colony throughout the period reported.

### Pathology

Each mouse found dead, or euthanized for humane reasons, was fixed as soon as possible. Incisions were made in the cranium, thorax, and abdomen, and the specimen then immersed in 10% neutral buffered formalin for storage at room temperature. Random samples from each of the three sites were later sent to the University of Washington Geropathology Research Program for gross inspection and for preparation of slides for histological examination. During gross examination and dissection, all obvious external and internal abnormalities were recorded. Tissues evaluated and collected for histologically examination included: decalcified cross section of skull with brain (when intact); lung; mediastinal lymph node and fat; thymus (when able to identify); heart; kidney; adrenal glands; liver; spleen; pancreas; mesenteric fat and lymph node(s); thyroid gland; salivary glands; reproductive tract (consisting of uterus and ovaries for female mice and testis, epididymis and seminal vesicles for male mice); and any additional lesions noted grossly. The gastrointestinal tract was inspected grossly but not routinely examined histologically due to autolysis. Tissues were routinely processed and paraffin embedded, after which 4-5 micron sections were stained with hematoxylin and eosin (H/E) for histological assessment.

Tissues were examined for a single major abnormality such as a malignant tumor extensively involving an organ or involving multiple organs; severe inflammatory disease; or an end-stage degenerative process such as cardiomyopathy or glomerulonephropathy. If such a process was identified, then this was considered the cause of death (COD). In some mice, more than one severe abnormality was identified, and in these cases death was attributed to two or more distinct lesions; these are listed in the results as “multiple” COD cases. If the carcass was in generally good postmortem condition but no obvious cause of death was identified, the COD was listed as “unknown.” If autolysis precluded critical histological evaluation of tissues for cause of death, a designation of “autolysis” was made. Some incidental lesions were recorded as present or absent, and other incidental lesions were graded with a severity score.

### Hemoglobin A1C (HbA1C)

Male and female mice at age 7 months were fed control or Cana diets for 7 to 8 months. These mice were not involved in the lifespan protocol. Blood was collected into EDTA-coated tubes from mice that were fasted for 6 hours starting at 8-9 AM. Samples were shipped to the University of Texas Health Science Center and HbA1C percentage for each whole blood sample was measured by immunoassay using a DCA Vantage Analyzer (Siemens Healthineers, USA) according to manufacturer’s instruction.

### Measurement of canagliflozin

Canagliflozin concentrations were measured in mouse serum, as well as in food pellets, using a validated bioanalytical assay with HPLC and tandem mass spectrometric detection (details available on request). The assessment of food levels included measures of within-pellet and between-pellet homogeneity and drug stability. Mean levels in prepared food averaged 85% of expected (nominal) concentration, with a range of 53% -180% of expected over 11 batches. Within- and between-pellet heterogeneity averaged 8%.

### Statistical Analysis

The analytical plan was specified prior to the initiation of the study. Survival was evaluated using the log-rank test, stratified by site, conducted separately for males and females. P-values are reported without adjustment for multiple comparisons. Secondary analyses of survival were conducted on data from each of the three test sites. The Wang/Allison (WA) test (24) was used to assess an analogue of “maximal lifespan;” this method uses the Fisher Exact test to contrast the proportions of mice alive or dead at the 90^th^ percentile age of the joint (shared) survival table for Control and Cana mice together. The primary test employed the sum of live and dead mice taken from each test site, separately for each sex, using the 90^th^ percentile age for that site and sex. Drug effects on body weight were assessed by the Student’s t-test, separately at each age for each sex, pooling across sites. Drug effects on specific inferred cause(s) of death, and on lesions scored as present or absent, were evaluated using the Fisher Exact test, pooling across site and sex. Drug effects on lesions that were graded were assessed using the t-test. HbA1C results were analyzed by two-way ANOVA, with Sex and Drug (Cana) as the predictor variables.

## Results

### Survival

In the context of the NIA Interventions Testing Program (ITP), groups of genetically heterogeneous (UM-HET3) mice born in 2016 were placed on a chow diet containing Cana at 180 parts per million (ppm) at the age of 7 months and followed for survival. There were approximately 50 males and 50 treated females at each of the three test sites (TJL = the Jackson Laboratory; UM = University of Michigan; and UT = University of Texas Health Science Center at San Antonio). Each site enrolled approximately 100 male and 100 female control mice, which also served as the control group for the other drugs tested in 2016. Date of death was recorded, and mice euthanized for humane reasons were considered to have died on the date of euthanasia. **Figure 1** shows the survival curves for each combination of sex and site, as well as the survival data pooled across all three sites. **Table 1** collects summary statistics for each site and for the pool. The pre-specified primary outcome was the log-rank test for the pooled data set, calculated separately for each sex, and stratified for site within sex.

**Table 1:** Lifespan Statistics for Cana and Control Mice – Pooled and Site-Specific Results.

| Rx | Group | Sex | Count | Median (Days) | Percent Change in Median | p-value | 90th %ile (Days) | Percent Change in 90th %ile | WA p-value |
| --- | --- | --- | --- | --- | --- | --- | --- | --- | --- |
| Cont_16 | Pool | M | 303 | 787 |  |  | 1047 |  |  |
| Cana | Pool | M | 156 | 897 | 14 | < 0.0001 | 1143 | 9 | < 0.0001 |
| Cont_16 | TJL | M | 102 | 749 |  |  | 1005 |  |  |
| Cana | TJL | M | 54 | 880 | 17 | 0.001 | 1159 | 15 | 0.003 |
| Cont_16 | UM | M | 102 | 814 |  |  | 1071 |  |  |
| Cana | UM | M | 51 | 946 | 16 | 0.001 | 1170 | 9 | 0.001 |
| Cont_16 | UT | M | 99 | 799 |  |  | 1064 |  |  |
| Cana | UT | M | 51 | 866 | 8 | 0.060 | 1099 | 3 | 0.39 |
| Cont_16 | Pool | F | 304 | 874 |  |  | 1050 |  |  |
| Cana | Pool | F | 136 | 886 | 1 | 0.512 | 1059 | 1 | 0.50 |
| Cont_16 | TJL | F | 96 | 883 |  |  | 1057 |  |  |
| Cana | TJL | F | 48 | 912 | 3 | 0.173 | 1097 | 4 | 0.26 |
| Cont_16 | UM | F | 92 | 858 |  |  | 1034 |  |  |
| Cana | UM | F | 44 | 897 | 4 | 0.263 | 1076 | 4 | 0.38 |
| Cont_16 | UT | F | 116 | 882 |  |  | 1058 |  |  |
| Cana | UT | F | 44 | 849 | -4 | 0.142 | 1003 | -5 | 0.56 |

**Figure 1:**
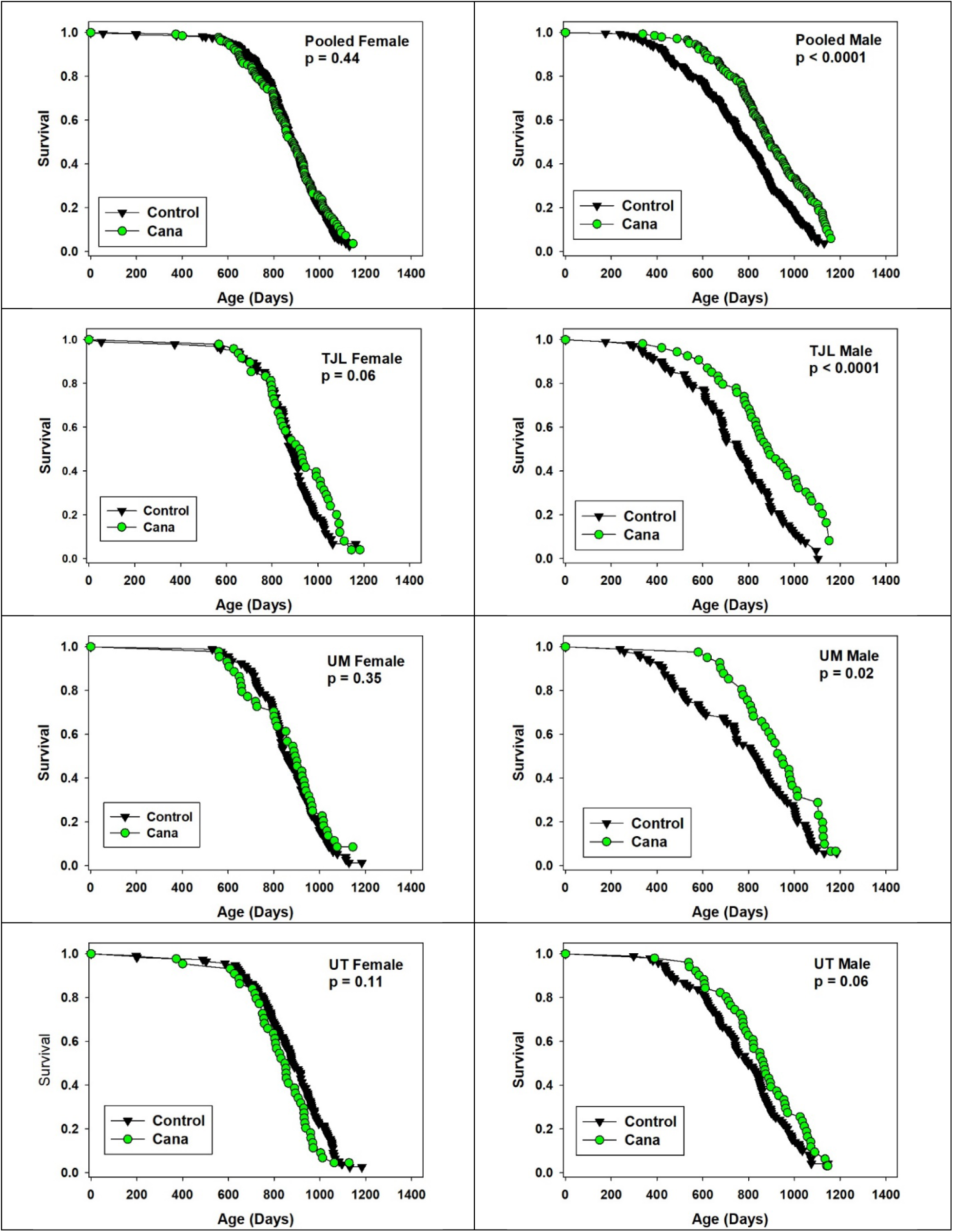
Survival curves for Cana mice and sex-matched Controls. The top row shows data pooled across sites, and the other three rows show site-specific results. Left panels are females and right panels are males. p-values show results of log-rank tests, not adjusted for multiple comparisons, with site as covariate for the pooled datasets (top row). There were no remaining live mice at the time of analysis.

For males, Cana increased median survival age by 14%; the log-rank test yielded p < 0.001. The log-rank test was also calculated for each site separately, and despite much lower statistical power, survival was improved at each site, with p < 0.001 at TJL and UM and p = 0.06 at UT. Age at the 90^th^ percentile survival point was taken as a pre-specified index of long-term (“maximal”) survival, and this increased by 9% in the pooled males, with site-specific values of 15%, 9%, and 3% for TJL, UM, and UT respectively. The Fisher Exact Test version of the Wang-Allison test (24) was used to evaluate the likelihood that the drug modified the proportion of survivors at the 90^th^ percentile; the calculated p-value, p < 0.001, suggests that Cana increases maximum survival in male mice. The WA statistic was significant at TJL and UM, but not at UT. None of these indices of improved survival were noted in female mice, for which the median survival age increased by only 1% (site-specific changes of 3%, 4%, and -4%), and age at the 90^th^ percentile survival was increased by a mere 1% (not significant).

Concentrations of Cana in plasma of young adult females ranged from 2.12 to 3.53 μg/ml (N = 4) after 8 weeks on Cana chow. Plasma of males ranged from 0.36 to 0.86 μg/ml (N = 3). This sex effect was significant at p = 0.003. Both sets of values are in the range of those reported for humans administered this drug (25). Thus the absence of a survival benefit in Cana females cannot be attributed to lower circulating levels of the drug. Husbandry technicians noted much higher urine output in the Cana mice of both sexes throughout the study.

### Weight

Cana blocks weight gain in both sexes, as shown in **Figure 2**. In females the effect on weight is dramatic (17% - 19%) and statistically significant (p < 0.0001) at each age after initiation of drug treatment at 7 months. Cana males are 5% to 8% lighter than controls at ages 12 and 18 months (p < 0.0002). Mean weight of Control males declines between 18 and 24 months, presumably related to onset of late-life illness and/or death of the heaviest males, but this effect is blunted in Cana mice. The male-specific alteration in lifespan is apparently not attributable to a male-specific blockage of midlife weight gain.

**Figure 2:**
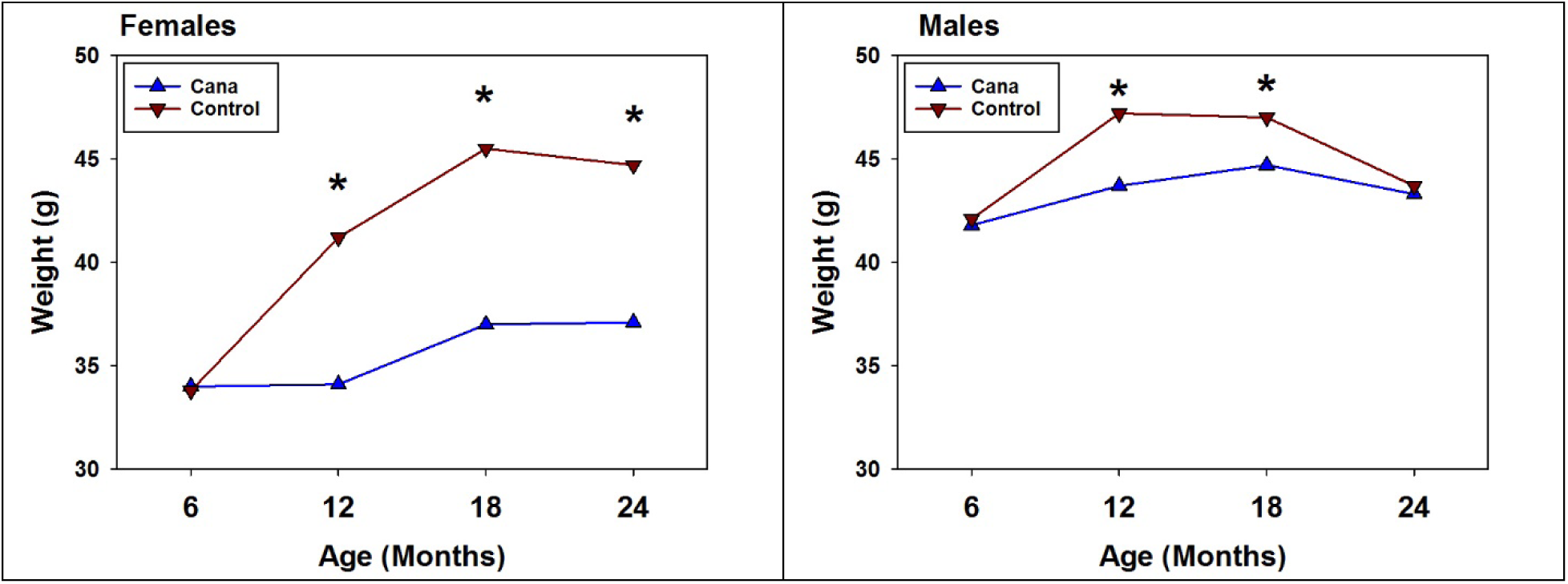
Mean weights for Cana and Control mice, ages 6 – 24 months. Values are mean levels for live mice, separately for each sex, pooled across sites. Cana treatment was initiated at 7 months. SEM values (not shown) ranged from 0.3 to 0.7 grams, and (*) p < 0.0002 for drug effect at ages 12 – 24 for females and 12 – 18 for males. N declined with age from 151 to 116 for Cana males and from 136 – 107 for Cana females, with approximately 2-fold higher numbers of Control mice.

### Hemoglobin A1c ((HbA1c) levels

HbA1c was evaluated in whole blood of 48 female and 45 male mice aged 7 months, after exposure to Cana for 7 to 8 months, with samples taken at all three test sites. There was no effect of Cana on HbA1c levels (Control males 3.8%, Cana males 3.7; Control females 3.5, Cana females 3.5), although the difference between sexes was significant at p < 0.0001 in a two-factor ANOVA. This result is consistent with our previous data on HbA1c in acarbose-treated mice (12), although Cana does diminish HbA1c in human diabetic patients (19).

### Cause of death

All mice dying at ages above 600 days were fixed, and a random sample (10 males and 10 females from each site in each of two drug conditions) were sent for evaluation to the University of Washington Geropathology Research Program without information as to Cana status. The mean age of Cana mice was 873 days (males) or 892 days (females), and the values for Control mice were 845 and 857 for males and females, respectively. Each case was evaluated by the same board-certified veterinary pathologist (JMS), who attempted to infer the cause(s) of death, i.e. the lesion(s) that appeared most likely to have led either to the death of the mouse or the decision to euthanize the mouse for humane reasons at the end of life.

**Table 2** shows the distribution of inferred causes of death (COD) among the mice. Statistical power is poor, because most individual diagnoses appeared in 7 or fewer of the 120 cases. The most common lethal lesion was hematopoietic neoplasia (HPN), a collection of hematopoietic neoplastic diseases, including lymphoma and histiocytic sarcoma, that are difficult to distinguish without special stains or molecular characterization. HPN was noted in 18/60 Cana mice and in 16/60 Controls. For most lesions, there were no effects of site (though as noted with very low statistical power), but HPN as a cause of death was more common at TJL (16/28 cases in which a COD could be inferred) than at UM and UT (18/70 non-autolyzed cases, p = 0.003). None of the classes of lethal illness shows a statistically significant difference between treated and control mice. For Cana mice, there were 45 cases for which a single cause of death could be assigned; this excludes the 10 cases where multiple lesions were deemed contributory and the 5 cases with advanced autolysis. Of these 45 mice, 34 (76%) died of some form of neoplastic disease. Similarly, 36 of the 44 evaluable Control cases (82%) died of neoplasia (p = 0.12 for contrast with Cana mice.)

**Table 2:** Inferred Causes of Death in Cana and Control Mice.

| Inferred Cause | Cases |  |  |
| --- | --- | --- | --- |
|  | Cana | Control | Total |
| Hematopoietic neoplasia | 18 | 16 | 34 |
| Hemangiosarcoma | 2 | 3 | 5 |
| Liver neoplasia | 3 | 4 | 7 |
| Lung carcinoma | 2 | 5 | 7 |
| Neoplastic, other | 9 | 8 | 17 |
| Atrial thrombosis | 5 | 2 | 7 |
| Non-neoplastic, other | 6 | 6 | 12 |
| Multiple or Unknown | 10 | 11 | 21 |
| Autolysis | 5 | 5 | 10 |
| Total | 60 | 60 | 120 |

Notes to Table 2. "Neoplastic, Other" category includes bone tumor (N=2), mammary adenocarcinoma (3), renal carcinoma (1), metastatic carcinoma of unknown primary site (4), unclassifiable neoplasia (6), and pituitary adenoma (1). "Non-neoplastic, other" includes endometritis (1), nephropathy (2), urinary tract obstruction (3), amyloidosis (2), and pyelonephritis (4).

### Incidental pathology at end of life

In addition to attempting to infer a likely cause of death, the pathologist recorded presence or absence of lesions in multiple tissues, and assigned a grade of severity for a separate set of lesions. **Supplemental Table 1** shows 21 lesions (in adrenal, atrium, lung, heart, liver, kidney, mammary gland, thalamus, pancreas, spleen, and thyroid) that were scored as present or absent, with p-values showing results of the Fisher Exact Test comparing Cana to Control mice. Drug treatment had no significant effects on frequency of any of these lesions. Supplemental Table 2 shows 14 lesions that were graded on scales of 0-2 or 0-4, depending on the lesion. The p-value shown reflects the two-sided Student t-test contrasting treated and control mice. Only one form of pathology met the criteria for statistical significance: the mean level of liver telangiectasia/angiectasis was 0.41 for Cana and 0.09 for Control mice (p = 0.01). Since this lesion is graded on a scale of 1 to 4, with a score of 1 corresponding to minimal change, it seems unlikely that this drug effect has any clinical implication. With that exception, the data suggest that Cana-treated mice show similar incidence and severity of most forms of end-of-life pathology.

## Discussion

Interventions, whether diets, mutations, or drugs, that slow aging and postpone multiple forms of late-life illness provide valuable tools for investigation of the cellular and molecular mechanisms for aging, as well as hints as to where those tools might most usefully be applied. Among the seven ITP-tested drugs that have shown significant longevity benefit, Cana, acarbose, 17aE2, and rapamycin have the largest effect on median lifespan, and each of these also has a significant effect on our measure of maximum lifespan (24), in one or both sexes. Surprisingly, acarbose, 17aE2, rapamycin can each produce significant lifespan extension even when started at ages 60% to 70% of the median survival age of controls, i.e. 16 to 20 months of age (10, 11) (and unpublished data for 17aE2). Lifespan studies in which Cana is initiated at older ages are now under way at each ITP site. Terminal (end of life) necropsies for each of these four drugs, including the data on Cana shown here, lead to the same conclusion: the spectrum of lethal and non-lethal late-life illnesses is not changed in quality or severity, even though the treated mice are older at time of death. Although this conclusion is limited by the low statistical power of small case series, the implication is that the drugs extend healthy lifespan, i.e. produce longer lifespan by delaying the onset or progression of the diseases most likely to lead to death or terminal morbidity, most of which are neoplastic disease in UM-HET3 and other mouse stocks. A comprehensive series of cross-sectional necropsies of rapamycin-treated mice (26), euthanized at 22 months of age, supported this inference by showing nine varieties of age-dependent lesions, in multiple organs, whose incidence was reduced by rapamycin.

The results of these studies show an unexpected degree of sexual dimorphism. The percentage benefit of Rapamycin, at each tested dose, is higher in females than in males, although this seems likely to reflect the higher blood rapamycin levels, in females, at matched doses in food (8). It is unclear whether one sex or the other would show preferential benefit at fixed blood levels of rapamycin. Acarbose, both in the original report and in an independent follow-up study (12, 15), led to significant lifespan extension in both sexes, but with much larger effects in males. The non-feminizing steroid 17aE2, in contrast, produced benefits only in males; 17aE2-treated males lived significantly longer than females whether or not the females had received this agent (11, 12). Two other tested drugs also produced significant lifespan benefits in male mice only. Nordihydroguiaretic acid (NDGA), an antioxidant with anti-inflammatory properties, led to male-specific lifespan increase of up to 12% (p = 0.006 by log-rank test), although without a significant increase in maximal lifespan; the benefit, and its limitation to males, was replicated in an independent study (11-13). Protandim, a mixture of botanical extracts known to activate the Nrf2 stress-response pathway, also led to increased lifespan in male mice only (11). Only one other tested agent, the amino acid glycine, produced significant lifespan extension in both male and female mice, and the effect, though seen at each of the three ITP test sites, was quite small, yielding less than a 5% increase in median longevity (14). Effects of acarbose, including an increase glucose tolerance and mTORC2 function (27) and a reduction of subscapular fat depots (15), were seen principally in males, and the effects of 17aE2 on metabolite profiles, glucose control, grip strength, and rotarod performance were also male-specific (28). Both acarbose and 17aE2 reduce mid-life hypothalamic inflammation in males only (17). More work will be needed to work out the molecular and physiological explanation(s) for these sex-specific effects, but it is notable that at least some of these effects are blocked by castration of young adult male mice prior to the initiation of treatment by acarbose or 17aE2 (27, 28), suggesting that levels of male hormones, even in post-pubertal adults, sensitize mice to lifespan extension by these drugs.

Cana and acarbose each reduce serum glucose in mice, or people, who have recently consumed a carbohydrate-rich meal, but they do so through partly different mechanisms. In addition to potently inhibiting SGLT2, Cana also reduces glucose uptake via SGLT1 in the small intestine and kidney, albeit with less potency than its effect on SGLT2 (29, 30). Physiologically, Cana suppresses glucose reuptake in the kidney, as well as delaying glucose absorption in the small intestine (18). Acarbose inhibits breakdown of polysaccharides to absorbable monosaccharides in the intestine. These observations thus suggest strongly that the lifespan benefits of each drug, and presumably the other benefits already shown in acarbose-treated mice, are a consequence of lower maximal post-prandial glucose levels. Metabolically, Cana and other SGLT2 inhibitors (SGLT2i) lower body weight, HbA1c, and peak glucose levels in humans; improve insulin resistance; increase serum ketones; enhance fatty acid oxidation; and elevate beta-cell function (18). Physiological effects in mice are not yet well delineated, and it is noteworthy that neither Cana nor acarbose, at the doses used by the ITP, seems to diminish HbA1c, suggesting that the lifespan and health benefits in mice may depend more on peak glucose levels than on average glucose levels integrated over the day. In our studies the dramatic effects on weight shown in both males and females in Figure 2 show that Cana has strong effects on fuel use and metabolic partitioning in both sexes, but additional studies will be needed to evaluate sex-specific modulations of relevant factors, particularly including insulin production and sensitivity, fat cell homeostasis and adipokine production, and hypothalamic and gastrointestinal circuits that modulate appetite and satiety. Notably, no major sex-specific effects of Cana have been noted in clinical trials documenting its beneficial effects in humans. However, our results are consistent with prior studies of long-term Cana administration in rats (31). In these studies, Cana, in a dose-dependent manner, extended rat lifespan, with a much greater effect observed in males than in females. As with other ITP drugs for which sexual dimorphic benefits are observed, the basis for the sex specificity of Cana on rodent lifespan remains unclear.

The end-of-life necropsy data for Cana mice are consistent with previous findings on mice whose lifespan was extended by 17aE2, acarbose, and rapamycin (9, 12). Specifically, the spectrum of lethal and non-lethal lesions, with a few exceptions, is similar between control and drug-treated mice, but delayed in timing. We see two complementary implications, one drawn from pathophysiology and the other from biogerontology. From the first perspective, our results justify follow-up studies to learn more about how these drug-induced changes, such as the blunting of peak glucose levels that accompany acarbose and Cana treatment, might delay or decelerate the most frequent forms of age-related illnesses, such as neoplasms of many tissues and the wide range of degenerative diseases seen in mice and humans, and can begin to test dietary and pharmacological interventions aimed at controlling glucose peaks. In parallel, researchers focused on questions in the basic biology of aging can use these drugs as experimental tools for testing hypotheses about how aging might be linked upstream to glucose or mTOR or steroid balance, and downstream to disability and dysfunction.

Cana and other SGLT2i show a wide spectrum of beneficial effects in humans and rodent models, which in principle could contribute to extension of mouse lifespan by Cana. The basis for these protective effects is not completely understood, and may not directly reflect the effects of Cana on glucose homeostasis. For example, Cana reduces systolic and diastolic blood pressure, fatal and non-fatal cardiovascular events, and hospitalization for heart failure (19). The potent cardioprotective effects of Cana and other SGLT2i are not observed to the same degree in patients treated with other, comparably potent, glucose-lowering agents, suggesting that pleiotropic effects of SGLT2i may be involved (20). Since most UM-HET3 mice die of neoplastic disease, the effects of Cana on cancer may be particularly relevant for its ability to extend mouse lifespan. In preclinical studies, Cana administered at high doses to rats increased the frequency of adrenal and renal tubular tumors, and induced Leydig cell hyperplasia and adenomas at all doses in males (31). In a recent meta-analysis, however, SGTL2i use in humans was not associated with an overall increased cancer risk, and Cana specifically was associated with a reduced risk of gastrointestinal cancer (32).

Hypothetically, Cana might reduce mortality associated with neoplasia in mice by modulating glucose metabolism in tumor cells. The metabolic requirements of rapidly dividing cancer cells are quite distinct from those of normal cells, most of which are ordinarily non-mitotic. In contrast, cancer cells must constantly generate large quantities of new biomolecules for macromolecular synthesis and cell replication. For most cancer types, glucose, and particularly glycolytic intermediates derived from it, represents a critically important fuel (33). SGLT1 and SGLT2 are functionally expressed on human prostate and pancreas cancers, and Cana treatment of a pancreas cancer xenograft potentiated the effects of genotoxic chemotherapy (34). Cana inhibits growth of SGLT2-expressing hepatocellular cancer xenografts via inhibition of glycolysis and suppression of tumor angiogenesis (35), and by reducing β-catenin expression (36). In a mouse model of nonalcoholic steatohepatitis-induced hepatocellular carcinoma, Cana reduced hepatic triglyceride accumulation, inflammation, and tumorigenesis (37). Cana has also been proposed to suppress growth of prostate and lung cancer cells via a non-SGLT1/2-dependent mechanism involving inhibition of mitochondrial Complex I (38). Thus Cana treatment may be acting, in part, by suppressing initiation and/or progression of age-associated cancers in UM-HET3 mice, through SGLT1/2-dependent or -independent mechanisms, an idea to be explored in follow-up studies.

Combining COD and incidental lesion scores is a powerful approach to evaluate the effects of a drug on progression of age-related pathology over a lifespan. Male mice given Cana showed an increase in lifespan compared to controls, but with similar COD and incidental lesion scores, suggesting a delay in major and incidental lesion pathology associated with aging. A cross-sectional study, now under way, will produce specimens for histopathology study at a defined age, 22 months, at which the complications of selective death and variable post-mortem interval have been greatly reduced. Such cross-sectional designs do not provide useful insights into possible drug effects on distribution of COD diagnoses, but can provide a more finely grained look at drug effects on age-dependent changes in multiple tissues and organs (26). COD and incidental lesion scores can be used as a framework for establishing pathological endpoints (39) and can be combined with physiological performance tests to further validate the effectiveness and safety of Cana or other drugs, either alone or in combination with other previously validated anti-aging drugs.

Prospective clinical trials for Cana effects in humans might be enriched by inclusion of endpoints that are signals of age-dependent diseases and physiological change that are nominally independent from the effects of the diseases, such as diabetes, for which the drug is prescribed. In parallel, secondary analysis of databases that include long-term follow-up on patients who are receiving Cana therapeutically might help to test the idea that this agent retards clinically important aspects of multi-system aging in men or women or both. The ITP results using both Cana and acarbose suggest that many aspects of age-related pathology, including those that lead to death in mice, could be sensitive to modulators of peak glucose levels, rather than integrated measures of average glucose values, with implications for both dietary and pharmacological avenues for prevention of late-life diseases in humans. Delineation of the mechanisms connecting Cana to slower aging, whether by effects on glucose or potentially through other pathways, are bound to have implications for our understanding of both mammalian aging and organ-specific pathophysiology.

## Acknowledgements

These studies were supported by NIH grants U01AG022303, R01AG05431, R01GM101171, P30AG013319, P30AG024824, U01AG022307, P30AG050886, U24AG066346, U01-AG-022308, and U24AG056053. We wish to thank Roxann Alonzo, Nelson Durgin, Ilkim Erturk, Melissa Han, Vicki Ingalls, Zhou Jiang, Jenna Klug, Pam Krason, Yuhong Liu, Natalie Perry, Wenbo Qi, Lori Roberts, and Jacob Sheets for expert technical assistance.

---

**Supplemental Table 1:**
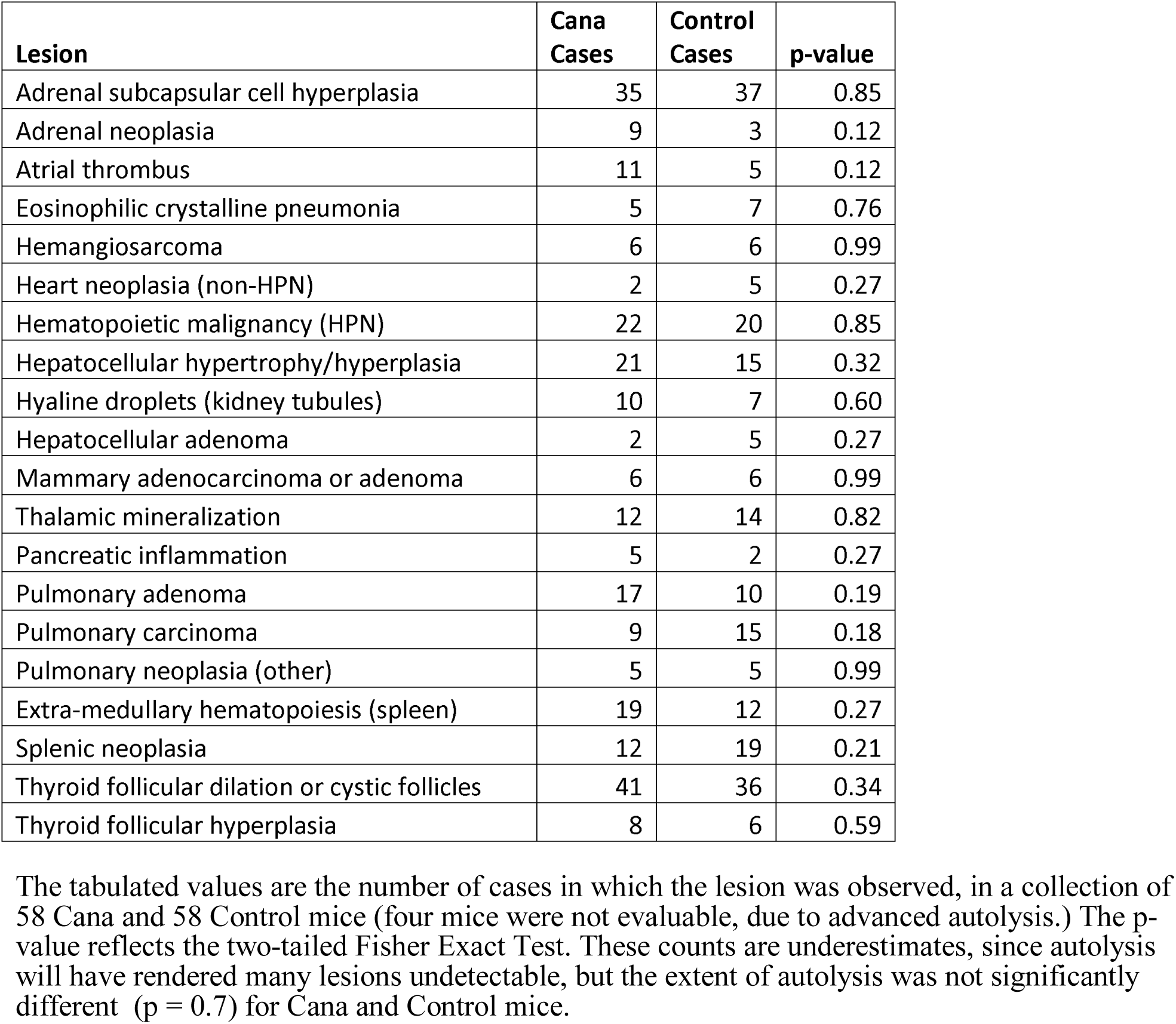
Incidental Pathology Findings – Lesion Counts.

| **Lesion** | **Cana Cases** | **Control Cases** | **p-value** |
| --- | --- | --- | --- |
| Adrenal subcapsular cell hyperplasia | 35 | 37 | 0.85 |
| Adrenal neoplasia | 9 | 3 | 0.12 |
| Atrial thrombus | 11 | 5 | 0.12 |
| Eosinophilic crystalline pneumonia | 5 | 7 | 0.76 |
| Hemangiosarcoma | 6 | 6 | 0.99 |
| Heart neoplasia (non-HPN) | 2 | 5 | 0.27 |
| Hematopoietic malignancy (HPN) | 22 | 20 | 0.85 |
| Hepatocellular hypertrophy/hyperplasia | 21 | 15 | 0.32 |
| Hyaline droplets (kidney tubules) | 10 | 7 | 0.60 |
| Hepatocellular adenoma | 2 | 5 | 0.27 |
| Mammary adenocarcinoma or adenoma | 6 | 6 | 0.99 |
| Thalamic mineralization | 12 | 14 | 0.82 |
| Pancreatic inflammation | 5 | 2 | 0.27 |
| Pulmonary adenoma | 17 | 10 | 0.19 |
| Pulmonary carcinoma | 9 | 15 | 0.18 |
| Pulmonary neoplasia (other) | 5 | 5 | 0.99 |
| Extra-medullary hematopoiesis (spleen) | 19 | 12 | 0.27 |
| Splenic neoplasia | 12 | 19 | 0.21 |
| Thyroid follicular dilation or cystic follicles | 41 | 36 | 0.34 |
| Thyroid follicular hyperplasia | 8 | 6 | 0.59 |

The tabulated values are the number of cases in which the lesion was observed, in a collection of 58 Cana and 58 Control mice (four mice were not evaluable, due to advanced autolysis.) The p-value reflects the two-tailed Fisher Exact Test. These counts are underestimates, since autolysis will have rendered many lesions undetectable, but the extent of autolysis was not significantly different ( $p = 0.7$ ) for Cana and Control mice.

**Supplemental Table 2:** Incidental Pathology Findings: Graded Lesions.

| Lesion | Cana | Control | p-value |
| --- | --- | --- | --- |
| Adrenal degeneration (*) | 1.76 | 1.90 | 0.55 |
| Cardiomyopathy | 1.46 | 1.65 | 0.26 |
| Glomerulonephropathy | 1.55 | 1.85 | 0.12 |
| Liver degeneration/necrosis | 1.06 | 0.64 | 0.10 |
| Liver telangiectasia/angiectasis | 0.41 | 0.09 | 0.01 |
| Macrovesicular hepatic lipidosis | 0.36 | 0.46 | 0.58 |
| Microvesicular hepatic lipidosis | 0.81 | 0.89 | 0.70 |
| Ovarian degeneration (*) | 3.67 | 3.42 | 0.26 |
| Pancreatic islet hyperplasia | 0.76 | 1.06 | 0.13 |
| Pancreatic exocrine atrophy | 1.09 | 0.76 | 0.39 |
| Salivary gland lymphoid aggregates | 1.29 | 1.30 | 0.99 |
| Testicular degeneration (*) | 1.48 | 1.83 | 0.20 |
| Uterine cystic endometrial hyperplasia | 2.33 | 2.10 | 0.41 |

Lesions were graded on scales (0 – 2 or 0 – 4, depending on lesion). Values shown are mean levels in Cana and Control mice, shown with the p-value from a two-sided Student's t-test (equal variance assumption). The number of scorable cases varied among lesions, because autolysis renders some lesions unscorable while allowing others to be assigned a grade.
\*Scoring for adrenal gland, ovary, and testes includes assessment of a number of age-related lesions in these organs, including adrenal pigment and cortical changes; ovarian atrophy, interstitial cell hyperplasia, and pigment; and testicular atrophy and mineralization

